# Prevalence of viral frequency-dependent infection in coastal marine prokaryotes revealed using monthly time series virome analysis

**DOI:** 10.1101/2021.09.23.461490

**Authors:** Kento Tominaga, Nana Ogawa-Haruki, Yosuke Nishimura, Hiroyasu Watai, Keigo Yamamoto, Hiroyuki Ogata, Takashi Yoshida

## Abstract

Viruses infecting marine prokaryotes have large impacts on the diversity and dynamics of their hosts. Model systems suggest viral infection is frequency-dependent and constrained by the virus-host encounter rate. However, it is unclear whether the frequency-dependent infection is pervasive among the abundant prokaryotic populations with different growth strategies (i.e. *r-*strategy and *K-*strategy). To address this question, we performed a comparison of prokaryotic and viral communities using 16S rRNA amplicon and virome sequencing based on samples collected monthly for two years at a Japanese coastal site, Osaka Bay. Concurrent seasonal shifts observed in prokaryotic and viral community dynamics indicated that abundances of viruses correlated with that of their predicted host phyla (or classes). Co-occurrence network analysis between abundant prokaryotes and viruses revealed 6 423 co-occurring pairs, suggesting a tight coupling of host and viral abundances and their “one to many” correspondence. Although dominant *K*-strategist like species, such as SAR11, showed few co-occurring viruses, a fast succession of their viruses suggests viruses infecting these populations changed continuously. Our results suggest the frequency-dependent viral infection prevailed in coastal marine prokaryotes regardless of host taxa and growth strategy.

## Introduction

Marine prokaryotes are ubiquitous in the ocean and play key roles in the global biogeochemical processes [1]. Most of observed species (>35,000 species-level operational taxonomic units [OTUs], based on 97% 16S rRNA sequence identity) fall into several major taxa (phyla or classes for Proteobacteria), such as α-Proteobacteria (e.g. SAR11), Bacteroidetes (e.g. *Flavobacteriaceae*), and Cyanobacteria (e.g. *Synechococcus* and *Prochlorococcus*) [2, 3]. Although individual species have distinct ecological niches, they are often classified into one of two growth strategists based on their potential growth rate and temporal dynamics: (i) *K-*strategist (slow-growing and persistently dominant, e.g. SAR11) and (ii) *r*-strategist (fast-growing and opportunistic, e.g. *Flavobacteriaceae*) [4]. However, recent high-frequency sampling schemes (e.g. daily) uncovered that species not recognized as *r*-strategists exhibit drastic fluctuations (e.g. Marine Group II euryarchaeota) [5, 6]. Further, finely resolved populations (genotypes or strains) within a species-level OTU often show distinct temporal dynamics [7–11], indicating species described as *K-*strategist can show frequent fluctuation.

Viruses infecting prokaryotes are abundantly present in the ocean and estimated to lyse 20–40% of the prokaryotic cells each day [4, 12, 13]. Viruses are thought to infect their specific hosts (often restricted to strains within a species) in a frequency-dependent manner, in which the encounter rate between the viruses and their hosts is a determinant for the infection rate [14, 15]. Thus, viruses infect host populations that become abundant and frequencies of host and viruses oscillate over time, leading to the maintenance of the diversity of the host community [16, 17]. Moreover, mathematical models have demonstrated that a prokaryotic species with faster growth rate can be susceptible to viral infection [17]. This trend allows *K-*strategists to reach a higher abundance than *r*-strategists because of their higher resistance against viral infection by cryptic escape through reduced cell size and/or specialized defense mechanisms [4, 18]. However, the discovery of SAR11 viruses questions this prediction [19]. It is currently unclear whether *K-strategists* suffer from viral infection or viral infection is prevalent in abundant prokaryotes regardless of their growth strategies..

Previous monthly observations of microbial communities have revealed that seasonal oceanographic features have a strong influence on the prokaryotic community [20, 21]. Seasonal variability of viral community also have been reported using PCR-based analysis [22, 23] and viral metagenomics (viromics) [24–26]. Although viruses are obligate parasites, viral seasonality was often discussed independently from the seasonality of their hosts except for few prokaryotic-virus pairs (e.g. *Synechococcus/Prochlorococcus*) [23, 27] because of the difficulty in connecting uncultured viruses and their hosts [13, 28].

In this study, we aimed to solve the two fundamental questions whether viral infection is prevalent among abundant prokaryotic populations or the way viruses infect differs depending on the taxa and/or growth strategies of their hosts. For this purpose, we monitored prokaryotic and viral communities at a eutrophic coastal site, Osaka Bay, monthly for two-years. We compare the community dynamics of viruses and that of their putative hosts using the *in silico* host prediction analysis [29, 30] and prevalence of viral infection is discussed based on the potential virus-host pairs determined through their co-occurrence dynamics.

## Materials and methods

### Sampling and processing

Seawater samples (4 l) were collected at a 5 m depth at the entrance of Osaka Bay (34°19′28″N, 135°7′15″E), Japan, within 3 h before or after high tide, between March 2015 and November 2016, at monthly intervals. Seawater was filtered through a 142 mm-diameter (3.0 μm pore size) polycarbonate membrane (Millipore, Billerica, MA) and then sequentially through 0.22 μm-pore Sterivex filtration units (SVGV010RS, EMD Millipore). After filtration, filtration units were directly stored at -80 °C for subsequent DNA extraction. The filtrates were stored at 4°C before treatments. Water temperature and salinity were monitored using fixed water intake systems of the Research Institute of Environment, Agriculture and Fisheries, Osaka prefecture. Nutrient concentrations (NO_3_-N, NO_2_-N, NH_4_-N, PO_4_-P, and SiO_2_-Si) were measured by continuous flow analysis (BL TEC K.K., Japan.).

### rRNA gene amplicon sequencing analysis

For prokaryotic community analysis, DNA was extracted from the stored filtration units as previously described [31, 32]. Total 16S rDNA was amplified using a primer set based on the V3–V4 hypervariable region of prokaryotic 16 S rRNA genes [33] with added overhang adapter sequences at each 5′ end according to the sample preparation guide (https://support.illumina.com/content/dam/illumina-support/documents/documentation/chemistry_documentation/16s/16s-metagenomic-library-prep-guide-15044223-b.pdf). Amplicons were sequenced using MiSeq sequencing system and MiSeq V3 (2 × 300 bp) reagent kits (Illumina, San Diego, CA).

Paired-end 16S rDNA amplicon sequences were merged using VSEARCH with the “-M 1000” option [34]. Merged reads containing ambiguous nucleotides (i.e., “N”) were discarded. The remaining merged reads were clustered using VSEARCH to form operational taxonomic units (OTUs) at a 99% sequence identity threshold. Singleton OTUs were discarded. The representative sequences of the remaining OTUs were searched against the SILVA ribosomal RNA gene database (release 138) [35] to taxonomically annotate OTUs using SINA [36] at a 99% sequence identity threshold. Abundant OTUs were defined as OTUs exceeding 1 % relative abundance by assuming the reported minimum host cell density for effective viral infection (≒10^4^ cells/ml) [37] and typical coastal marine prokaryotic cell density (≒10^6^ cells/ml) [38].

To identify statistically relevant variants within abundant OTUs, we applied minimum entropy decomposition (MED) [11] as previously reported [7]. All the sequences from each 99% OTU were aligned using MAFFT v7.123b (-retree 1 - maxiterate 0 -nofft -parttree) [39]. The alignment of sequences containing positions with entropy of >0.25 position was decomposed, and decomposition continued until all positions had entropy of <0.25. The minimum number of the most abundant sequence within each amplicon sequence variant (ASV) needed to exceed 50 and ASVs that did not exceed 1% of the parent OTU composition were discarded [7].

### Virome sequencing, assembly, classification, and calculation of relative abundance

The filtrate containing viruses was concentrated via FeCl_3_ precipitation [40] and purified using DNase and a CsCl density centrifugation step [41]. The DNA was then extracted as previously described [42]. We failed to obtain enough amount of DNA for virome sequencing for one sample (February 2016), the sample was removed from the analysis. Libraries were prepared using a Nextera XT DNA sample preparation kit (Illumina, San Diego, CA) according to the manufacturer’s protocol, using 0.25 ng viral DNA. Samples were sequenced using a MiSeq sequencing system and MiSeq V3 (2 × 300 bp) reagent kits (Illumina, San Diego, CA).

Viromes were individually assembled using SPAdes 3.9.1 with default *k*-mer lengths [43]. Additionally, we used scaffolds of these assemblies (hereafter referred to as contigs for simplicity). Circular contigs were determined as previously described [44]. Contig sequences were clustered at 95% global average nucleotide identity with cd-hit-est (options: -c 0.95 -G 1 -n 10 -mask NX, 549 redundant contigs were discarded) [45]. A total of 5 226 mts-OBV contigs (monthly time series Osaka Bay viral contigs, >10 kb, 62 — 926 contigs/samples, including 202 circular ones) were obtained. Genome completeness and quality of mts-OBV contigs were evaluated using checkV (v0.7.0) [46]

In addition, this assembly generated 181 131 short contigs (i.e., from 1 kb up to 10 kb). The abundance of these contigs was assessed based on the relative abundance of terminase large subunit genes (*terL*) as previously described [32]. In total, 4 666 genes were detected as putative *terL* genes (i.e., genes with the best hit to PF03354.14, PF04466.12, PF03237.14, and PF05876.11). Fragments per kilobase per mapped million reads (FPKM) for putative *terL* genes were calculated using in-house ruby scripts.

The mts-OBV contigs with complete viral genomic sequence set collected in a previous study [44] were used for viral abundance estimation based on read mapping. The complete viral genomic sequence belonged to one of the following two categories: (i) 1 811 environmental viral genomes (EVGs; all are circularly assembled genomes, 45 were assembled in Osaka Bay in a previous study [44]) derived from marine virome studies; (ii) 2 429 reference viral genomes (RVGs) of cultured dsDNA viruses. Genus-level genomic OTUs (gOTUs) were previously assigned for complete genomes based on genomic similarity score (S_G_) using ViPTree [47]. For the mts-OBV contigs, if a sequence showed a high similarity to one of the complete genomes (with S_G_ > 0.15), the sequence was assigned to the gOTU of the most similar genome as previously described [32, 44]. Quality controlled virome reads were obtained through quality control steps as previously described [44]. These reads were mapped against the viral genomic sequence set using Bowtie2 software with the “--score-min L,0,-0.3” parameter [48]. FPKM values were calculated using in-house ruby scripts.

### Viral host prediction

First, we assigned putative host groups based on the genomic similarity with viral genomic sequence set collected in a previous study [44]. If mts-OBV contigs were classified into the same gOTU with the viruses with a known (via cultivation) or predicted (by genomic content [44]) host group, the host group was assigned to the contigs. We also compared similarity with mts-OBV contigs, the viral genomes deposited in a virus-host database (as of October 2018), and recently reported isolates [49, 50].

In addition, for the viruses without assigned host groups via genomic similarity, we performed *in silico* host prediction based on the nucleotide sequence similarity between viruses and prokaryotes as previously described [30, 51, 52]. First, a total of 220 103 viral genomes/contigs derived from marine viromes were collected and used for the analysis [24, 44, 53–55] (**Supplementary Table 1**). For the putative host genomes, we collected a total of 8 016 MAGs/SAGs from marine metagenomic or single cell genomic studies [56–60]. From Pachiadaki *et al*, we only used 1 040 high quality SAG assemblies with ≥ 80% completion [60]. To remove the contamination of virus-like contigs from the MAGs/SAGs, 14 967 contigs classified as viral-like sequences using VirSorter (category 1, 2, and 3) [61] were discarded (**Supplementary Table 1**). Details of each prediction method were reviewed previously [29].

#### CRISPR-spacer matching

CRISPR-spacer sequences were predicted using the CRISPR Recognition Tool [62], and then a total of 13 305 sequences were extracted. Detected spacer sequences and spacer sequences deposited in CIRSPRdb [63] were queried against viral genomes using the BLASTn-short function [64] with the following parameters: At least 95% identity over the whole spacer length and only 1–2 SNPs at the 5′-end of the sequence was allowed.

#### tRNA matching

tRNAs were recovered from MAGs/SAGs and viral genomes using ARAGORN with the ‘-t’ option [65]. A total of 213 939 and 31 439 tRNAs were recovered from MAGs/SAGs and viral genomes, respectively. The recovered prokaryotic and viral tRNAs with 111 385 tRNAs deposited in GtRNAdb [66] were compared using BLASTn [64] and only a perfect match (100% length and 100% sequence identity) was considered as indicative of putative host-virus pairs.

#### Nucleotide sequence homology of prokaryotic and viral genomes

Viral genomes/contigs were queried against prokaryotic MAGs/SAGs and prokaryotic genomes in NCBI RefSeq (as of December 2019) using BLASTn [64]. Only the best hits above 80% of identity across alignment with a length of ≥1500 bp were considered as indicative of host-virus pairs. For the prediction based on MAG/SAGs contigs, we performed taxonomic validation of the matching contigs in MAG/SAGs as previously described [30]. Viruses belonging to the same gOTU were assigned consistent host groups according to a previous study [44], with three exceptional gOTUs (G404, G405, and G495), which annotated multiple host lineages. For the contigs assigned to the three gOTUs, genomic similarity among the same gOTU members were calculated and the potential host of each contig was assigned based on the most similar genomes/contigs which was annotated via host prediction

### Statistical analyses

Before statistical analyses, 16S rRNA amplicon reads were rarefied using the “vegan” package in R (20 803 reads per sample, based on minimum sample size) [67]. To examine within-sample alpha-diversity (Shannon diversity, evenness, and richness) and beta-diversity (Bray-Curtis similarity: 1 - Bray-Curtis dissimilarity, for all of the possible pairwise combinations among all of the sampling points), we used the vegan package in R [68]. Mantel tests were performed using R and the vegan package [68] only on fully overlapping sets of data. Pairwise correlations between estimated abundance of prokaryotic ASVs and viral contigs (having putative host information and exceeding FPKM >10 at least a month, 2 735 contigs) on fully overlapping sets of data were then determined via Spearman correlation (P<0.01, Q<0.05) as implemented in the local similarity analysis program. [69, 70]. Network visualizations of correlation matrices were generated using Cytoscape_v3.8.0 [71].

### Estimation of the growth strategy of ASVs

We established indexes for the approximation of the *r* (intrinsic rate of natural increase) and *K* (carrying capacity) of each ASV by monitoring their monthly dynamics. For the approximation of the *r* of each ASV, the maximum increase of the normalized relative rank (0-1) per month was applied. Similarly, for the approximation of *K* for each ASV, the length of continuously dominant month (>0.1% relative abundance, 1-18 months) of each ASV was applied.

### Detection of SNPs

Reads were mapped to the viral contigs using Bowtie2 with a “--score-min L,0,- 0.3” [48] and the resulting alignment files were converted to BAM format and sorted using samtools [72]. The average genome entropy of the contigs which exceeded more than 10 coverage each month was computed using the DiversiTools (http://josephhughes.github.io/DiversiTools/).

### Data availability

Sequences obtained from the observations were deposited at the DNA Data Bank of Japan (DDBJ) under project number PRJDB10879. Raw sequence reads can be found under accession numbers DRX260081 to DRX260115 and assemblies of viromes can be found under BioSample SAMD00279559.

## Results and discussion

### Overview of prokaryotic and viral communities in Osaka Bay

We obtained 2.8 M paired-end reads (24 168 to 846 565 reads per sample) from the 16S rRNA gene V3-V4 region amplicon sequencing libraries derived from 18 collected samples and these sequences were clustered into 35 191 OTUs (1 462 to 18 268 OTUs per month) with a sequence identity threshold of 99% (species-level populations, **Supplementary Table S2**). The prokaryotic community was dominated by α-Proteobacteria (41%), γ-Proteobacteria (21%), Bacteroidetes (19%), and Cyanobacteria (7%) at the phylum level (class level for Proteobacteria).

To explore viral community composition, we obtained 60 M paired-end reads of viromes (929 884 to 8 124 354 sequences per sample), which were generated from the virus size fraction of 17 samples that were concomitantly collected with the prokaryotic size fractions (**Supplementary Table S2**). After decontamination of prokaryotic sequences, 5 226 virus-like large contigs (> 10 kb, monthly time series Osaka Bay viral contigs: mts-OBV contigs) were obtained, including 202 circularly assembled viral genomes (**Supplementary Table S2**). In this study, we refer to these contigs operationally as species-level viral populations, according to the previous proposal in viral ecology [73]. The majority (∼75%) of mts-OBV contigs showed high genomic similarity (genomic similarity score; S_G_ > 0.15; see [44] for the definition of S_G_) with one of the previously reported viral complete genomes [44] and the 202 circular genomes assembled in this study. Based on the S_G,_ these mts-OBV contigs were classified into 314 gOTUs (**Supplementary Table 2)**. On average, 40% of virome reads (29 to 53% per sample) were mapped on the mts-OBV contigs or previously reported viral genomes [44]. The mts-OBV contigs occupied 96% relative abundance on average for individual samples (based on the FPKM values calculated from read counts). Relative abundance of terminase large subunit genes (*terL*) of the whole set of contigs (>1 kb) indicates that all mts-OBV contigs (>10 kb) were ranked at the top (>30%) of the whole community in at least one sample (the lowest of maximum relative abundance was 0.0115%, 16Jan_NODE_472, **Supplementary Figure S1**).

Alpha-diversity (Shannon index) of the viral community was significantly higher than that of the prokaryotic community (*p<* 0.001, **Supplementary Figures S2A-B**). Both richness and evenness were also significantly higher in the viral community than in the prokaryotic community (**Supplementary Figure S2C-F**, *p<* 0.001). It should be noted that prokaryotic diversity was evaluated via single marker gene analysis (i.e., 16S rRNA) but viral diversity was evaluated via whole genome sequencing. Thus, the methodological difference could have caused the relatively higher diversity of the viral community. Another possible explanation for the higher viral diversity is that a prokaryotic species can be infected by more than one viral species at each time point (discussed below).

### Seasonal dynamics of prokaryotic and viral communities

We investigated seasonal dynamics of prokaryotic and viral communities using the Bray-Curtis similarity index between all possible pairs of samples (136 pairs, 1- to 17-month intervals). Both prokaryotic and viral communities showed clear seasonal patterns, with a peak of average similarity at an interval of about 12 months, representing the same seasons, and the bottom of average similarity at an interval of 6 months, representing opposite seasons (**Figure 1**). Prokaryotic community dynamics were concordant with seasonal environmental variables, such as water temperature and inorganic nutrients, which increased in summer (June to September) presumably because of the increasing river inflow during the rainy season (**Supplementary Table S3, Supplementary Figure S3)**. The similarity between samples was systematically lower for the viral community than that for the prokaryotic community (**Figure 1**, discussed below). The viral community composition was significantly correlated with the prokaryotic community composition, as well as the seasonal environmental variables (Mantel *rho* = 0.504, *p* < 0.01, **Supplementally Table S3**).

**Figure 1.**
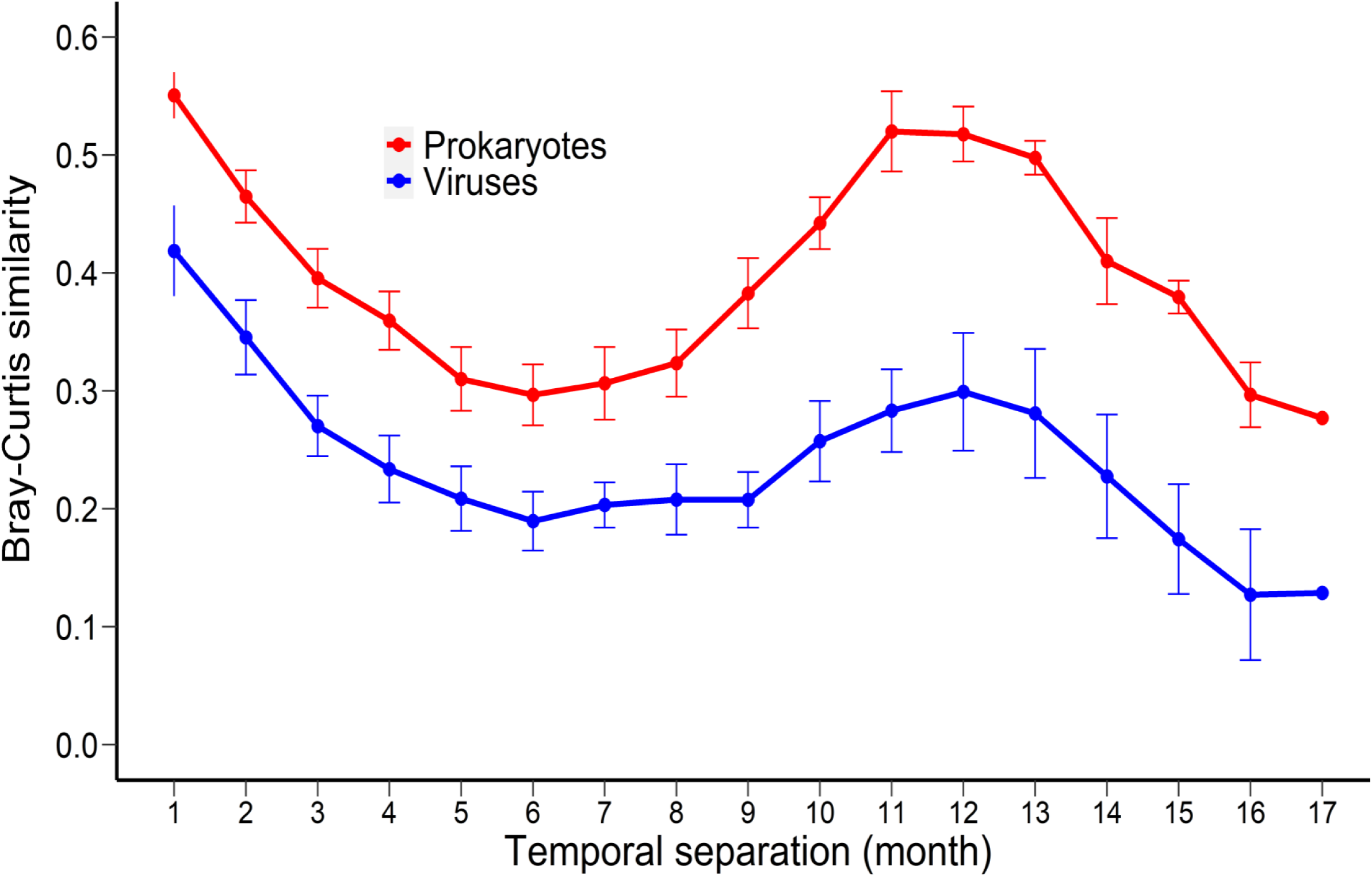
Seasonality of the prokaryotes and viruses at the Osaka Bay (OB) during observation. The Bray-Curtis community similarity index was calculated among all of the possible sample pairs from normalized abundances of prokaryotic OTUs and OBV contigs and plotted as a function of the number of months separating their sampling.

Given that each virus can only propagate in its specific host, and thereby the viral community composition is shaped by prokaryotic community composition, abundance of each virus might reflect the abundance of its host. To test this hypothesis, compositions of prokaryotic and viral communities were compared using the information of predicted viral hosts (mostly host phylum- or class level composition). Putative host groups of viruses were predicted using four commonly used genome-based *in silico* prediction methods (similarity with known viruses, CRISPR-spacer match, tRNA match, and genome homology). First, based on the similarity with cultured viruses, putative host groups of 951 mts-OBV contigs (22 gOTUs) were predicted (*Synechococcus/Prochlorococcus*, 182 contigs; SAR11, 501 contigs; SAR116, 214 contigs; *Roseobacter*, 31 contigs; others, 23 contigs, **Supplementally Table S4**). Similarly, putative host groups of 504 mts-OBV contigs (39 gOTUs) were predicted based on the similarity with uncultured viral genomes considering previous assignment of putative hosts (Bacteroidetes, 468 contigs; MGII, 36 contigs [30, 44], **Supplementally Table S4**). For other 1 460 mts-OBV contigs (α-Proteobacteria, 35 gOTUs, 621 contigs; Bacteroidetes, 80 contigs; γ-Proteobacteria, 236 contigs; δ-Proteobacteria 326 contigs; others, 53 contigs, **Supplementally Table S4-5**), putative host groups were predicted via the sequence similarity (i.e. CRISPR-spacer matching, tRNA matching, and genome homology) between viral (mts-OBVs with previously reported >200,000 marine viral genomes [24, 44, 53–55]) and prokaryotic genomic data sets (>8 000 marine prokaryotic metagenome-assembled genomes in previous studies [56–60] and the genomes in the NCBI RefSeq database). Altogether, we assigned potential host groups for 2 844 mts-OBV contigs (α-Proteobacteria, 1 375 contigs; Bacteroidetes, 548 contigs; δ-Proteobacteria, 326 contigs; γ-Proteobacteria, 250 contigs; Cyanobacteria 190 contigs, **Supplementally Table 4**).

Major phyla (or classes for Proteobacteria) in the prokaryotic community did not change drastically but the relative abundance of several phyla (classes) exhibited remarkable seasonal dynamics (**Figure 2**). The seasonal dynamics of the predicted viral hosts resembled the seasonal dynamics of prokaryotes (**Figure 2**). For example, Cyanobacteria (79% of reads were assigned to OTU_8, *Synechococcus*) dominated in summer (up to 9.6% and 22.6% of the community in June 2015 and July 2016, respectively, **Figure 2**) and *Synechococcus* virus abundance also increased in summer (up to 5.3 and 12.1% of the community in August 2015 and August 2016, respectively, **Figure 2**). Similarly, the relative abundance of Bacteroidetes increased from winter to spring (up to 33.7% of the community in May 2016, **Figure 2**) and Bacteroidetes virus abundance also increased during spring (up to 30.2% of the community in May 2016, **Figure 2**). Relative abundances of both SAR11 (from 5 to 47% of the community, **Figure 2**) and SAR11 viruses (from 9 to 22% of the community, **Figure 2**) showed changes over time but they were always abundant throughout the observed period. Therefore, virally community appear to generally follow the dynamics of their host.

**Figure 2.**
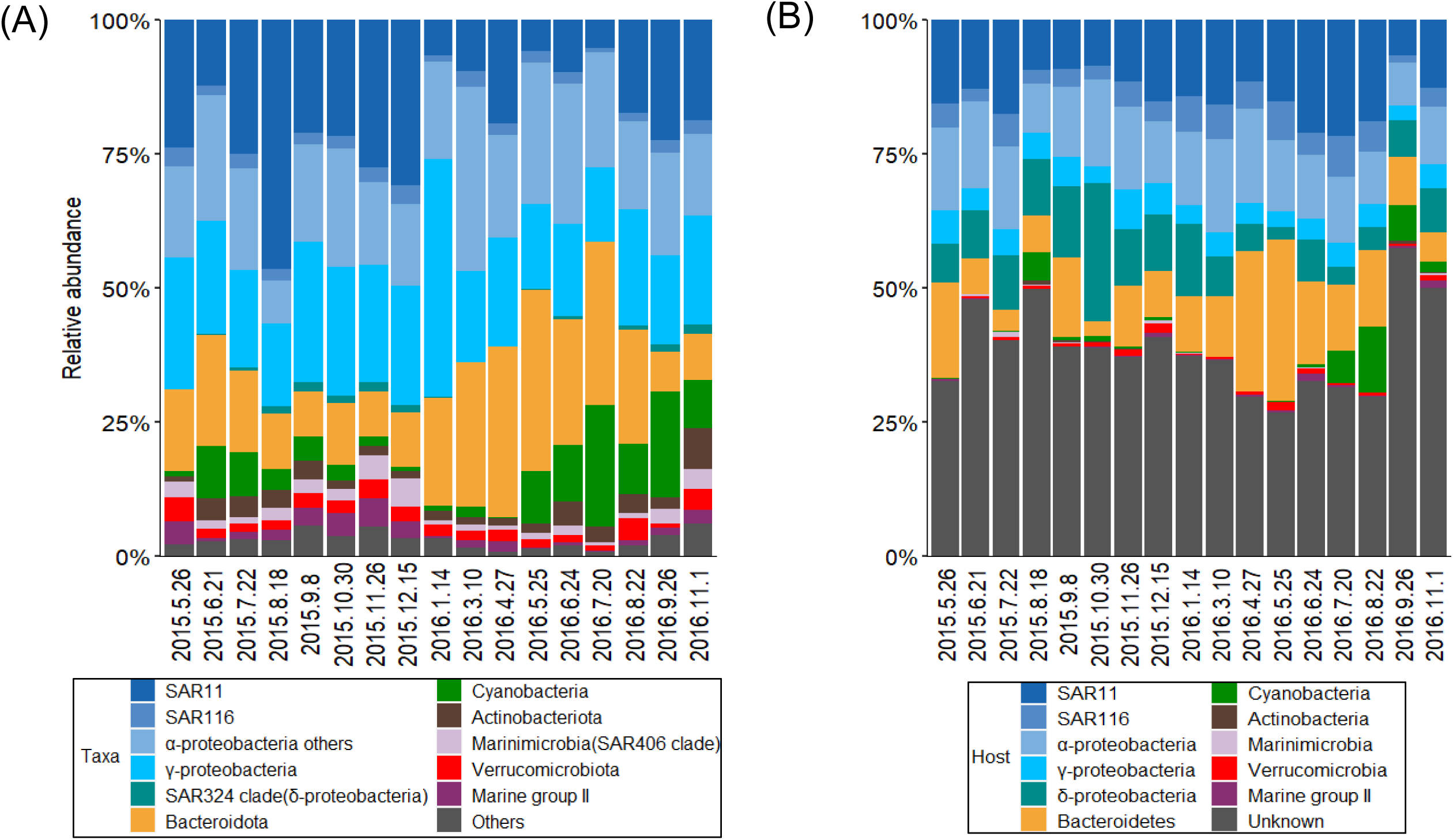
Comparison of prokaryotic and viral taxonomic community composition based on the host prediction. (A) Relative abundance of phylogenetic groups of prokaryotic communities. Quality-controlled reads were clustered into OTUs with sequence identity of 99% using VSEARCH (Rognes et al., 2016). These OTUs were classified at the phylum level (class level for Proteobacteria) using SINA (Pruesse et al., 2012). (B) Relative abundance of viruses based on their putative hosts assigned by host prediction. Normalized abundances of viral contigs were calculated from fragments per kilobase of per million reads mapped (FPKM) value.

However, viral abundance did not always match with their putative host abundance (**Supplementally Figure S4**). For example, the proportion of putative γ-Proteobacteria viruses was lower compared with that of γ-Proteobacteria and the proportion of putative δ-Proteobacteria viruses was much higher compared with that of δ-Proteobacteria (**Figure 2**). The lack of a tight correlation between viral and host abundance may not be surprising. The host prediction based on genome analysis in this study was mostly at the phylum or class level except for contigs showing similarity with cultured viruses, such as *Synechococcus/Prochlorococcus* cyanoviruses, while typical prokaryotic viruses could only infect specific host species or strains. Further, although our analysis annotated putative hosts at nearly 60% of the viral community, remaining populations without host prediction may lead to the underestimation of viruses infecting some taxa. The difference in burst sizes among viruses, which have been estimated to range from 6 to 300 in the marine environment [74], can also influence the estimation of viral abundance. Next, to investigate whether viral abundance increased according to specific host abundance, we statistically examined associations (i.e. co-occurrence) between the viruses and ASVs extracted from the abundant 73 prokaryotic OTUs.

### Co-occurrence network analysis between the abundant prokaryotes and viruses

To examine the dynamics of closely related (nearly strain-level) variants within each OTU, 114 ASVs (1∼4 ASVs per OTU, **Supplementally Figure S5**) were extracted from the abundant 74 OTUs via minimum entropy decomposition [7, 10, 11]. Then, pairwise correlations (co-occurrence network) between the 114 prokaryotic ASVs and the viral species, which were predicted to infect the prokaryotic ASVs via host prediction (e.g. 37 Bacteroidetes ASVs and 548 mts-OBV contigs predicted as Bacteroidetes virus), were determined via Spearman’s correlations. In total, 6 423 significant correlations between 104 prokaryotic ASVs and 1 366 viral species were detected (**Figure 3, Supplementary Figure S6**). The majority (88.6%) of prokaryotic ASVs correlated with at least one viral species. In contrast, only 34% and 31% of prokaryotic ASVs positively and negatively correlated with environmental variables, respectively (Spearman correlations (*r*>|0.6|, P<0.01, Q<0.05, **Supplementary Table S6**). The number of co-occurring viral species ranged from 0 (13 ASVs) to 359 (ASV6-1, classified into *Planktomarina*) and the median value was 16.

**Figure 3.**
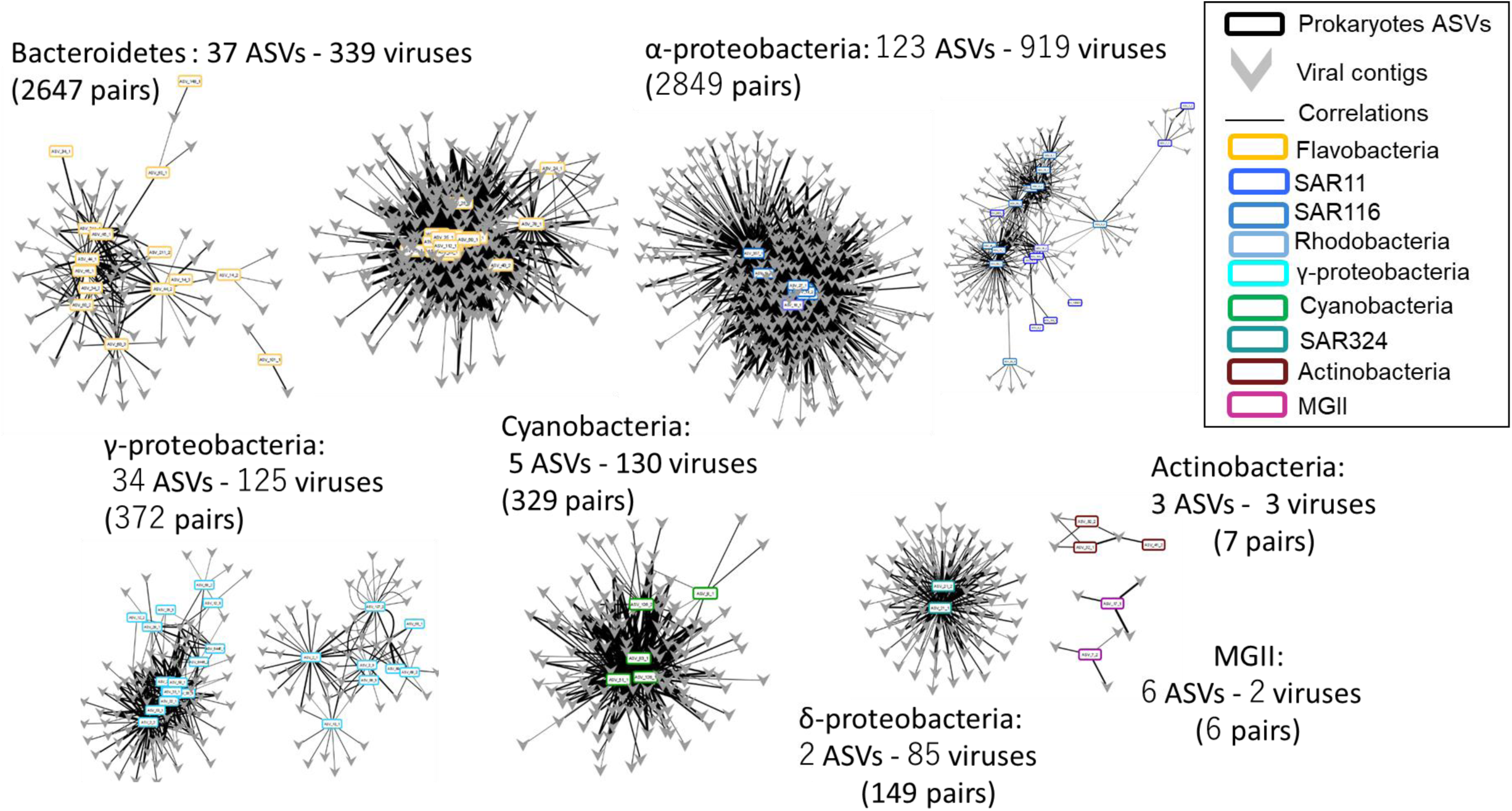
Broad overview of detected positive correlations between prokaryotic ASVs and viral populations which potentially infect each prokaryotic taxa based on host prediction analysis. (A) Flavobacteria and their viruses. (B) α-proteobacteria and viruses. (C) γ-proteobacteria and their viruses. (D) Cyanobacteria and their viruses. (E) Other major groups (SAR324, Marine group II, and Actinobacteria) and their viruses. Prokaryotic nodes are circles and viral node are v-shapes. Node color indicates prokaryotic taxa. Solid lines are positive correlations.

Using the detected 6 423 putative virus-host pairs, we examined whether the viruses were abundant when their putative host was abundant. First, four cyanobacterial ASVs and co-occurring 130 cyanovirus species were examined. Since substantial numbers of *Synechococcus*/*Prochlorococcus*-virus pairs have been reported in culture-based studies [75–78], host prediction for cyanoviruses is likely to be reliable. These cyanoviral species were more dominated in the viral community when their co-occurring ASVs exceeded predicted minimum host cell density for effective propagation of prokaryotic viruses (10^3^ cells/ml [79] or 10^4^ cells/ml [37], **Figure 4, Supplementary Figure S7, S8**). Thus, cyanobacterial viral species were not abundant or often undetectable when their putative hosts were less abundant, but they became dominant when putative host abundance increased. This viral increase with host abundance was also observed in 98 other prokaryotic ASVs and their co-occurring viral species (**Figure 4, Supplementary Figure S7, S8**). This result clearly indicates that frequency-dependent viral infection is prevalent in abundant prokaryotes at least between the detected virus-host pairs.

**Figure 4.**
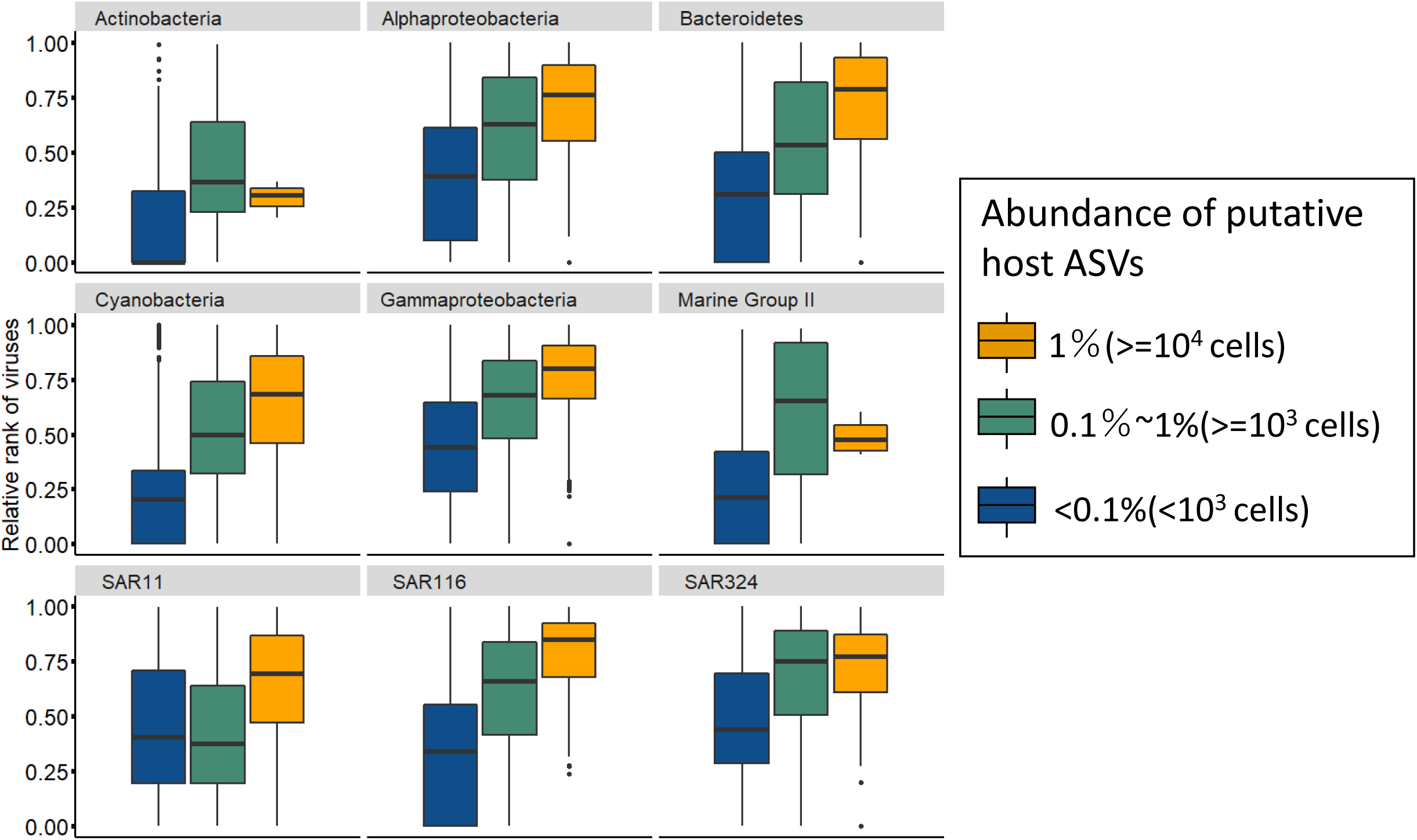
Increase of viral abundance according to the host cell density between co-occurring host-virus pairs. Normalized relative rank of each virus in community (0 ∼1) were plotted when their putative host relative abundance exceeding 1% (≒10^4^ cells/ml, yellow), 0.1% (≒10^3^ cells/ml, green), and below 0.1% (blue). Boxplots are constructed with the upper and lower lines corresponding to the 25th and 75th percentiles; outliers are displayed as points.

### Characterization of the virus-host interaction by host taxa

The community of viruses showed a higher alpha-diversity that the community of prokaryotes (**Supplementary Figure S2**), and the co-occurrence analysis indicated one-to-many associations between the host and viral populations (median 16 viral species per a prokaryotic ASV). This suggests that one abundant prokaryotic ASV can interact with multiple viral species. Note that the numbers of co-occurring viral species were overestimated since each contig could be a partial genome fragment derived from the same viral genome (average completeness of mts-OBV contigs was 39%, **Supplementary Table S4**). However, the contigs classified into different genera (average 8 gOTUs) often co-occurred with an ASV. Next, we characterized the “one to many” virus-host interaction network (i.e. how many viruses co-occurred with each ASV) with respect to their host taxa and host growth strategy.

The number of co-occurring viral species for prokaryotic ASVs was generally dependent on the predicted number of their viruses determined via host prediction (**Supplementary Figure S9**). For example, Bacteroidetes viruses (548 viruses) were the second most frequently observed ones and an average of 71.5 viruses co-occurred with Bacteroidetes ASVs (1–208 viruses per ASV, between 37 Bacteroidetes ASVs and 339 Bacteroidetes viruses). The number of co-occurring viruses could be overestimated because of the double count of co-occurring viruses between two co-occurring ASVs (if ASV-A and ASV-B co-occurred, the viruses co-occurring with ASV-A also can be included in the viruses co-occurring with ASV-B and vice versa. In fact, up to 16 ASV-ASV co-occurring pairs were detected for Bacteroidetes). In contrast, the taxa with less frequently detected viruses (e.g. MGII, 38 viruses) had a smaller number of co-occurring populations (0–3 viruses per ASV, **Supplementary Figure S9**). Thus, the number of co-occurring viral species might be underestimated in these taxa because of host prediction limitations. Exceptionally, SAR11 had relatively few co-occurring viral species even though there were more than 500 putative SAR11 viral species (**Supplementary Figure S9**). SAR11 is often regarded as a *K*-strategist, which is believed to be resistant to viral infection [4], and the growth strategy may influence the co-occurrence dynamics with viruses. Next, we examined the number of co-occurring viruses among prokaryotic ASVs classified in the same taxa depending on the growth strategy to solve this issue.

### Characterization of the virus-host interaction by host growth strategy

The growth strategy (*r* or *K*) of each prokaryotic ASV was defined by the indexes that we introduced (see methods). According to these, 13 ASVs were determined as *K*-strategist-like ASVs (i.e. *K*-index>12, *r*-index< 0.1). Among the 13 ASVs, seven were classified into SAR11 (**Supplementally Figure S10**). Twenty two of 57 ASVs belonging to the taxa previously predicted as *r*-strategist (i.e. *Flavobacteriaceae, Rhodobacteraceae, Vibrio*, and Marine Group II) were classified into the *r*-strategist-like ASVs (*K*-index<3, *r*-index>0.5, total 33 ASVs) (**Supplementally Figure S10**). Generally, *r-*strategist-like ASVs, such as members of Bacteroidetes, showed a large number of co-occurring viral species (**Supplementally Figure S10**). In contrast, *K-*strategist-like ASVs of *Synechococcus* and SAR11 showed relatively few co-occurring viral species (**Supplementally Figure S10**). The most abundant ASV of *Synechococcus* (ASV8-1, making up 76.7% of the whole cyanobacterial reads) and SAR11 (ASV1-1, occupied 7-64% of whole SAR11 reads of each month) showed 7 and 16 co-occurring viruses, respectively, even though 183 cyanoviruses and 500 SAR11 viruses were detected during the observation **(Supplementally Figure S10)**.

If a temporal switch of virus-host pairs occurred, co-occurrence analysis may fail to detect virus-host associations.. Therefore, we compared dynamics of the two dominant prokaryotic ASVs and viral species that did not co-occur with their predicted hosts. Representative sequence of ASV8-1 matched with the members of *Synechococcus* subcluster 5.1a at 100% of identity. Among the 53 cyanoviral species that did not co-occur with any cyanobacterial ASV, 41 species were classified into two gOTUs (G14, T7-like cyanosiphovirus, and G386, T4-like cyanomyovirus), which are known to infect subcluster 5.1a (e.g. *Synechococcus* sp. WH 8103, clade II), suggesting plausible interaction between ASV8-1 and these viruses. ASV8-1 especially dominated during summer (maximum 8% and 21% of prokaryotic community in June 2015 and July 2016, respectively, **Figure 5A**). Of these 53 viral species, abundances of which also increased in summer, four were only abundant in 2015 (from five to > 170 times abundant in 2015 than 2016) and other 38 species were more abundant in 2016 (from five to >300 times more abundant in 2016 than 2015) **(Figure 5A)**. Similarly, ASV1-1 of SAR11 was always abundant (**Figure 5B**) and SAR11 viruses occupied a major fraction of the viral community. However, abundant members of SAR11 viruses (309 contigs) were replaced in a relatively short time (a few months) (**Figure 5B**). These results suggest that the host-virus interaction might have been underestimated in the co-occurrence analysis and *K*-strategists also interact with multiple viruses based on their cell density.

**Figure 5.**
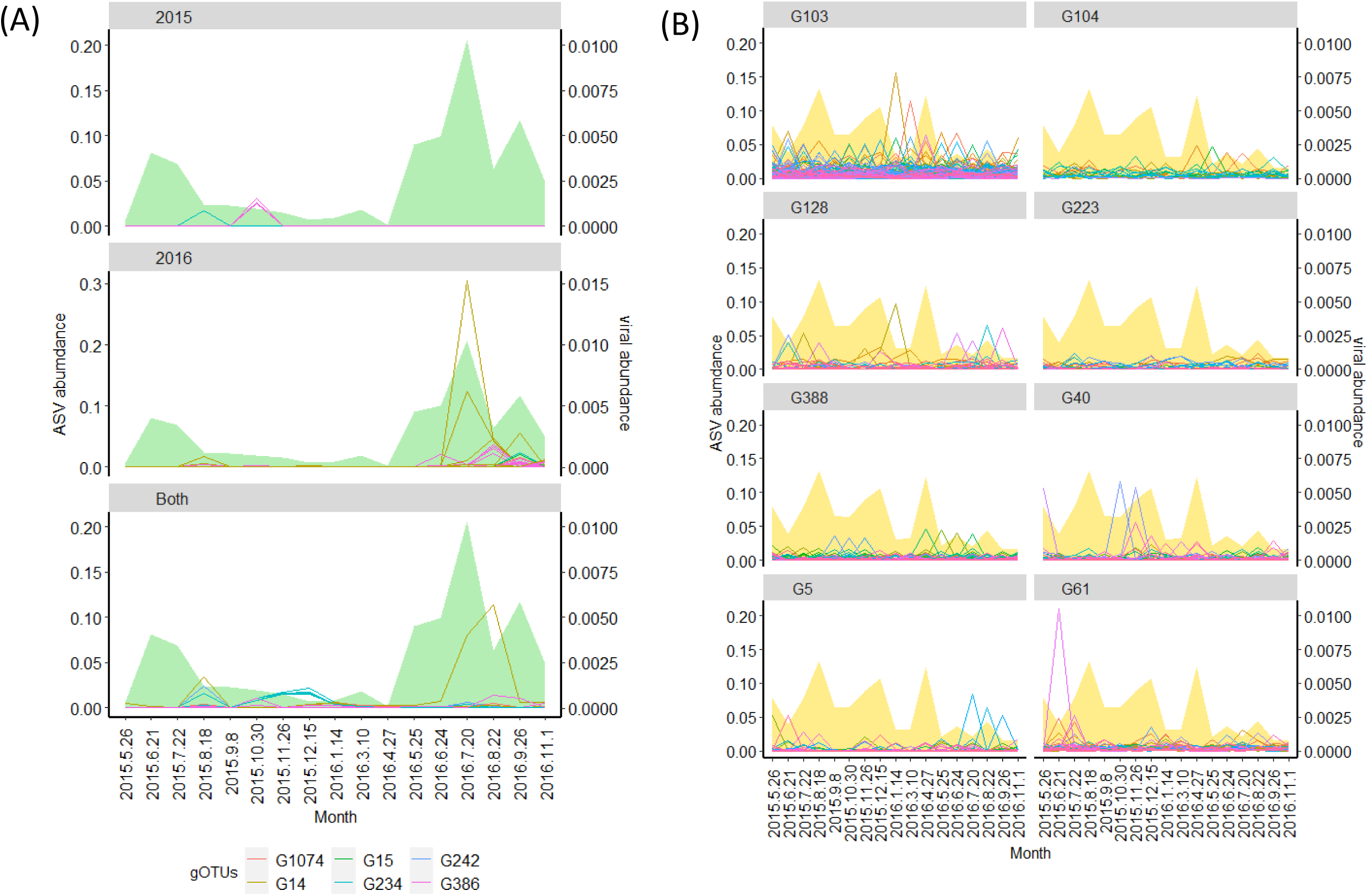
Dynamics of the most dominant prokaryotic population (ASV1-1 and ASV8-1) with viruses which predicted to infect these host taxa by host prediction analysis but did not co-occurred with any ASV. (A) Dynamics of ASV8-1 which classified into *Synechococcus* and 53 cyanoviruses which did not co-occurred with cyanobacterial ASVs. (B) Area chart represents relative abundance of the ASV8-1 and lines represents viral contigs over time. The panels were separated by viral annual pattern (2015 type, 2016 type, and both years, if the virus was more than five times abundant in one year comparing with another year, the virus was defined as year-specific virus). Colors represent gOTU (genus) of the virus. (B) Dynamics of ASV1-1 which classified into SAR11 clade and 309 putative SAR11 viruses which did not co-occurred with any SAR11 ASVs. Area chart represents relative abundance of the ASV1-1 and lines represents viral contigs over time. The panel were separated based on the classified gOTUs of each virus.

Finally, we investigated whether the observed viruses, including those not statistically detected as co-occurring viruses with hosts (e.g., 53 cyanoviruses and 309 SAR11 viruses in Figure 5), were also produced via increased contact frequency with hosts. To infer the contact frequency, we focused on single-nucleotide polymorphisms (SNPs) in viral genomes. SNPs of closely related viral populations were previously observed in abundant viral populations, such as freshwater cyanoviruses [80] and marine viruses in other coastal areas [24]. Since a recent study suggested that the majority of viruses observed in the virome were produced via diel and local viral-host interactions [32], it likely indicates that multiple infection events may lead to the generation of mutations through DNA replications. We thus hypothesized a frequent reproduction and mutations for abundant viruses with an increased contact frequency with their hosts. Therefore, SNPs from mts-OBV contigs with more than ten coverages (2 356 contigs) were calculated. We observed an increase of intrapopulation genetic diversity (SNPs quantified by average genomic entropy) as a function of overall population abundance regardless of their host taxa (**Supplementary Figure S11**). This result corroborates the notion that contact-rate is the key parameter for the viral reproduction regardless of whether they show a long term co-occurrence pattern with their hosts.

### Ecological interpretation inferred from virus-host dynamics

There are at least three possible mechanisms of the above-mentioned virus-host pair switch (**Figure 5**). First, more closely related prokaryotic populations that cannot be differentiated by the 16S rRNA gene polymorphism could co-occur with viruses. Previous studies focusing on the polymorphism of ITS sequences (ITS-ASV) in SAR11 and Cyanobacteria reported that ITS-ASV dynamics correlate more with viral dynamics, inferred from T4-like viral marker genes, than 16S-ASV dynamics of these taxa [7, 27]. Therefore, dynamics of more highly resolved populations (e.g. ITS-ASVs or whole genome sequence based-populations) might have synchronized with observed viral dynamics. Second, the temporal acquisition of host resistance or viral counter-resistance as often observed in culture model systems [83] may cause a switch of the dominant viral species. Third, it can be interpreted as a result of the founder effect, following host fluctuation via genetic drift [81]. Seasonal fluctuating of host population cause bottleneck effect, and therefore, the founder effect following the bottleneck effect enables the abundance of several viral species to equally increase. This was suggested as a mechanism of an incomplete selective sweep in the freshwater Cyanobacteria populations having different CRISPR-spacer genotypes [82]. The scenario is more plausible between ASV8-1 and their viruses because ASV8-1 experienced clear seasonal fluctuation (**Figure 5A**).

Altogether, we revealed that the frequency-dependent infection occurred in abundant prokaryotic populations according to the cell density via “one to many” host-virus correspondences regardless of the host growth strategy. One to many host-virus correspondences may suggest a prokaryotic species attacked by multiple viruses having a different infection strategy (e.g. different cell surface targets). This can cause difficulties in establish complete resistance toward multiple co-existing viruses and sustain continuous virus-host interaction in the environment. The difficulty of the emergence of “virus-free” species may be a potential mechanism for the prevailed frequency-dependent selection of abundant marine prokaryotes.

## Conclusion

Comparison of monthly dynamics between prokaryotic and viral communities indicated concurrent seasonal shifts at the whole community level. Concurrent seasonal shifts were also broadly observed between the corresponding virus and host pairs at the phylum or class level based on the host prediction analysis. We further statistically confirmed their co-occurrence via network analysis among abundant prokaryotic populations and their viruses regardless of the host taxa or growth strategies. These results suggested that abundant prokaryotes were exposed to frequent viral infection regardless of their taxa or growth strategy. It indicates that lysis of the abundant prokaryotes via viral infection have a considerable contribution to the biogeochemical cycling and maintenance of prokaryotic community diversity. Further, these abundant prokaryotic populations should reflect actively growing members of the community since they became dominant even though they suffered frequent loss by viral lysis.

## Acknowledgments

KT would like to thank Ryoma Kamikawa, Shigeki Sawayama, and Daichi Morimoto at the Kyoto University for technical comments and preparation of this manuscript. KT would also like to thank Hisashi Endo, Florian Prodinger, Hiroaki Takebe, Kentaro Fujiwara, and Tatsuhiro Isozaki at Kyoto University and Keizo Nagasaki at Kochi University for useful discussions. Computational work was supported by the Super Computer System, Institute for Chemical Research, Kyoto University. This study was supported by Grants-in Aids for Scientific Research KAKENHI (No. 17H03850 and No. 21H05057), and Challenging Exploratory Research (No. 26660171) from the Japan Society for the Promotion of Science (JSPS), Canon Foundation (No. 203143100025), JSPS Scientific Research on Innovative Areas (No. 16H06437), and the Bilateral Open Partnership Joint Research Project (Japan-Lithuania Research Cooperative Program) “Research on prediction of environmental change in Baltic Sea based on comprehensive metagenomic analysis of microbial viruses”.

## Supplementary Figure/Tables

**Supplementary Figure S1. Virome abundance of OBV long contigs as assessed by putative *terL* genes**.

Abundance of 1,078 mts-OBV long contigs (indicated by red) was assessed by the abundance of putative *terL* genes (from 4,666 contigs in total). y-axis represents the *terL* FPKM of each virus. Contigs (x-axis) are lined in order of the assembled month (from 2015 May to 2016 Nov).

**Supplementary Figure S2. Alpha diversity profiles of prokaryotic and viral communities in Osaka bay during observation**.

Average of Shannon H’ (A), richness (number of OTUs or contigs, C), and evenness (Pielou’s j: Shannon diversity divided by log richness, E) were calculated from normalized abundances of prokaryotic OTUs based on rarefied reads and viral contigs from fragments per kilobase of per million reads mapped (FPKM) value. The boxes represent the first quartile, median, and third quartile. Asterisks denote significance (Student’s t-test adjusted by Bonferroni correction., ***p< 0.001). The change of Shannon H’ (B), richness (D), and evenness (F) of prokaryotic and viral communities of the time-series were plotted.

**Supplementary Figure S3. Changes in environmental parameters at the Osaka Bay (OB)**.

Heatmap represents z-score transformed value of measured environmental parameters.

**Supplementary Figure S4. Relationship of relative abundance of prokaryotic taxa and viruses predicted to infect the corresponding prokaryotic taxa**.

x-axis indicate relative abundance of viruses at each month. y-axis indicate relative abundance of prokaryotes at corresponding month. Pro indicate the prokaryotic taxa and Vir indicate putative host of the viruses.

**Supplementary Figure S5. Dynamics of abundant prokaryotic OTUs and its decomposed ASVs**.

The yellow area-graph represents the relative abundance over time of each abundant OTU as a proportion of the whole community. The colored lines are the estimated relative abundance of each ASV (only >0.1% in abundance among whole community are shown) as a proportion of the whole community of prokaryotic sequences.

**Supplementary Figure S6. Dynamics of ASVs and their co-occurring viruses**

The yellow area-graph represents the normalized relative abundance (0 to 1) over time of each ASV. The dashed lines represents the normalized relative abundance (0 to 1) over time of each viruses which co-occurred with the ASVs. Only up to top 30 most abundant co-occurred viruses were show for each ASV.

**Supplementary Figure S7. Plots of relative abundance of co-occurring host-virus pairs**.

Relative abundance of each prokaryotic ASV and their co-occurring viruses at each month were shown. Black and red dot-lines represents 10^3^ cells/ml and 10^4^ cells/ml of host abundance, respectively.

**Supplementary Figure S8. Comparison of relative rank of viruses and host ASVs abundance among co-occurring host-virus pairs**.

Normalized relative rank of each virus in community (0 ∼1) were plotted when their putative host ASV relative abundance exceeding 1% (≒10^4^ cells/ml, yellow), 0.1% (≒ 10^3^ cells/ml, green), and below 0.1% (blue). Boxplots are constructed with the upper and lower lines corresponding to the 25th and 75th percentiles; outliers are displayed as points.

**Supplementary Figure S9. Number of virus-host co-occurring pairs by taxa**. Number of detected viruses by host prediction of each host taxa were shown as blue (first y-axis) and number of co-occurring viruses per an ASV (on average) by host taxa were show as yellow (second y-axis).

**Supplementary Figure S10. Distribution of the number of co-occurring viruses among prokaryotic ASVs based on their growth strategy inferred from approximated index of carrying capacity (K) and intrinsic rate of natural increase (r) based on their dynamics**.

x-axis indicates approximation index of *r* and y-axis indicates approximation index of *K*. Size of the circles represents the number of co-occurring viruses with each ASV. Color of the circles indicate the taxa of each ASV.

**Figure S11. Correlation of genome average entropy and abundance of OBV contigs calculated from SNP profiles**.

The graphs show the average genomic entropy of mts-OBV contigs and read coverage of the mts-OBV contigs at given time-series samples. The panel were separated based on the predicted hosts of the mts-OBV contigs.

**Supplementary Table S1. Basic statistics of microbial and viral genomes used for the host prediction analysis**.

**Supplementary Table S2. 16S rRNA amplicon and virome read sequences in each time series samples**.

**Supplementary Table S3. Rho values of Partial Mantel tests for prokaryotic and viral communities and environmental parameters**. The value in each box is the Rho value and data with *p* < 0.005 are indicated with * and with *p*<0.01 are indicated with **

**Supplementary Table S4. General genomic features and putative hosts of 5**,**226 mts-OBV contigs**.

**Supplementary Table S5. Putative virus-host pairs predicted in this study by methods based on CRISPR-spacers, tRNA, and host-virus genomic similarity**.

**Supplementary Table S6. Detected significant Spearman’s correlations (r>**|**0.6**|, ***p*<0.01**,***q*<0.05) between environmental variables and dynamics of ASVs**.

